# Systemic and Local Delivery of siRNA to the CNS and Periphery via Anti-IGF1R Antibody Conjugation

**DOI:** 10.64898/2026.01.30.702950

**Authors:** Mengnan Tian, Mehran Nikan, Miran Yoo, Stephanie Klein, Seung-Hwan Kwon, John Matson, Dongin Kim, Jinwon Jung, Sumin Hyeon, Byeong Min Yoo, Hyeon Ji Park, Michael Tanowitz, Asa Wahlander, Weon-Kyoo You, Hakju Kwon, Janel Huffman, Thazha P. Prakash, Sang Hoon Lee, Hien Zhao, Sungwon An

**Affiliations:** Ionis Pharmaceuticals Inc., 2855 Gazelle Ct, Carlsbad, CA 92010, USA; ABL Bio Inc., 456, Bongeunsa-ro, Gangnam-gu, Seoul, Republic of Korea

**Author notes:** To whom correspondence should be addressed. Correspondence may also be addressed to Hien Zhao. Equal contributions (Mengnan Tian, Mehran Nikan).

## Abstract

siRNA delivery platforms capable of accessing both central and peripheral tissues are critically needed to expand the therapeutic potential of oligonucleotides. To address this, we developed a novel siRNA-antibody conjugate by attaching an *Hprt*-targeting siRNA to an engineered antibody shuttle. This shuttle targets the insulin-like growth factor 1 receptor (IGF1R) using a fused antibody fragment (Clone F) and utilizes an antibody backbone with no tissue-relevant binding in this study. The resulting conjugate, designated Clone F-*Hprt*, demonstrated robust in vivo knockdown across multiple tissues. Clone F-*Hprt* demonstrated enhanced penetration into central nervous system (CNS) tissues compared to unconjugated siRNA following intracerebroventricular (ICV) and intravenous (IV) administration. In peripheral tissues, Clone F-*Hprt* achieved widespread knockdown in muscle, heart, and lung, consistent with IGF1R expression. The conjugate was well tolerated across all routes, including with repeated dosing. Although several receptor-mediated approaches for CNS delivery are progressing to the clinic (e.g., targeting the transferrin receptor), clinical validation remains to be demonstrated. Our findings highlight IGF1R as an alternative receptor capable of supporting delivery across both central and peripheral tissues, offering a complementary strategy for expanding the therapeutic landscape of oligonucleotide delivery.

## INTRODUCTION

Short-interfering RNA (siRNA)-based therapeutics are a rapidly evolving class of medicine that modulates the genetic causes of diseases in the clinic. By recruiting specific gene sequences to the RNA-induced silencing complex (RISC), siRNA leads to targeted gene silencing (1,2). Early siRNA therapeutic development faced challenges, such as poor tissue delivery and low stability. However, significant advancements in siRNA design, stability, and delivery methods have resulted in seven Food and Drug Administration (FDA)-approved siRNA therapies (3). A significant breakthrough in tissue-specific delivery came with the development of *N*-acetylgalactosamine (GalNAc), a ligand targeting liver-specific asialoglycoprotein receptors. This approach enabled efficient siRNA delivery to the liver, resulting in effective knockdown of genes expressed in hepatocytes (4). The success of GalNAc-mediated hepatic delivery has led to four FDA-approved siRNA drugs for liver-related disorders (1,5). Efforts to expand siRNA delivery beyond the liver are ongoing. For instance, lipophilic conjugates, such as C16 (2′-*O*-hexadecyl), have shown promise in delivering siRNAs to various brain tissues and the spinal cord in rodents and non-human primates (6). These lipophilic modifications enhance the ability of siRNAs to cross biological barriers and improve their cellular uptake. However, due to the tightly regulated barriers of the central nervous system (CNS), siRNA therapeutics targeting the brain often still require direct injection into the cerebrospinal fluid (CSF) for optimal efficacy (7). Antibodies have been widely developed as therapeutics for various indications, including cancers, autoimmune diseases, and neurodegenerative disorders, due to their specific binding capabilities. Beyond their use as single agents, antibodies have been explored as carriers for other therapeutic modalities. Examples include antibody-drug conjugates for cancer treatment and antibodies conjugated with neuroprotective peptides or enzymes for CNS-related pathologies (8-11). Despite this versatility, *in vivo* studies using antibody-siRNA conjugates remain limited, particularly in the context of tissue-specific delivery. For example, anti-transferrin receptor (TfR) antibody-siRNA conjugates have been evaluated by two groups, primarily for gene modulation in muscle tissue (12,13).

One promising strategy for CNS delivery involves the use of antibodies as molecular shuttles that exploit receptor-mediated transcytosis (RMT) across the blood-brain barrier (BBB). This mechanism utilizes antibodies that bind to transporters expressed on the surface of brain endothelial cells, allowing cargo to traverse the BBB and reach the parenchyma. Several groups have demonstrated successful CNS delivery of antisense oligonucleotides using antibody-based approaches targeting receptors such as TfR (12,13). However, the specific application of antibody-siRNA conjugates for CNS-targeted gene modulation remains at a very early stage and is yet to be systematically explored. In line with this strategy, a previous study demonstrated that an IGF1R-targeting scFv, Grabody B, functions as an effective molecular shuttle, mediating increased brain and CSF penetration of a therapeutic antibody (14). Building on this proof-of-concept, we utilized a related single-chain variable fragment (scFv) against IGF1R, designated Clone F, as the specific molecular shuttle for the siRNA conjugate in the current work.

To evaluate the potential of this shuttle system for oligonucleotide delivery, we aimed to investigate the impact of an engineered antibody shuttle targeting IGF1R on the biodistribution of oligonucleotides in both CNS and peripheral tissues by conjugating it with an siRNA targeting hypoxanthine-guanine phosphoribosyl transferase (*Hprt*) mRNA, a ubiquitously expressed housekeeping gene involved in purine metabolism. Surprisingly, the resulting Clone F-*Hprt* siRNA conjugate (Clone F-*Hprt*) significantly increased the potency of the unconjugated parent siRNA in the brain after intracerebroventricular (ICV) administration, despite its substantially larger size compared to the siRNA molecule. Furthermore, Clone F-*Hprt* effectively delivered *Hprt* siRNA into the brain following intravenous (IV) administration, resulting in reduced *Hprt* mRNA expression across multiple brain regions. Remarkably, it also achieved greater knockdown efficacy than unconjugated siRNA in peripheral tissues, including the lungs, heart, muscle, and liver following systemic dosing. These results open new possibilities for the delivery of various therapeutic modalities, including siRNAs, to the CNS, and indicate that a shuttle incorporating the Clone F scFv is applicable to tissue-targeted delivery of siRNAs with potentially superior cell penetration in both CNS and peripheral organs.

## MATERIALS AND METHODS

### siRNA synthesis

The antisense strand of *Hprt* siRNA was synthesized on NittoPhaseHL UnyLinker® solid support (317 μmol/g) using an AKTA Oligopilot 10 synthesizer at a 40 μmol scale, while the sense strand was synthesized on a commercially available 3′-Amino-Modifier C7 CPG solid support (Glen Research) to introduce a primary amine at the 3′-end. All phosphoramidites, including those for 2′-*O*-methyl and 2′-fluoro nucleosides, were dissolved at 0.1 M in acetonitrile, and the 5’-POM-(*E*)-Vinylphosphonate (VP)-2’-MOE-T phosphoramidite was used at 0.15 M in a 50:50 (v/v) mixture of toluene and acetonitrile. Each synthesis cycle began with detritylation using 15% dichloroacetic acid in toluene, followed by coupling with 1 M 4,5-dicyanoimidazole and 0.1 M N-methylimidazole in acetonitrile for 12 min per step, capping with 20% methylimidazole in acetonitrile (Cap A) and 20% acetic anhydride/30% 2,6-lutidine in acetonitrile (Cap B), and oxidation with 0.05 M iodine in 9:1 pyridine:water. Phosphorothioate linkages were introduced by treatment with 0.1 M xanthane hydride in 3:2 pyridine:acetonitrile. After synthesis, oligonucleotides were cleaved from the solid support, deprotected, and purified as reported previously.(15) Briefly, deprotection was performed by incubation in 10% diethylamine in 9:1 ammonium hydroxide:ethanol for 36 h at room temperature. The crude products were purified by ion-pair reversed-phase HPLC on an XBridge Prep C18 column using an acetonitrile gradient in 5 mM tetrabutylammonium acetate, followed by strong anion-exchange chromatography on SOURCE 30Q resin with 100 mM ammonium acetate and 1.5 M NaBr in 100 mM ammonium acetate, both in 3:7 acetonitrile:water. Final desalting was performed on a C18 reverse-phase column, and the purified oligonucleotides were dried in a vacuum concentrator. The *Hprt* siRNA duplex consists of an antisense strand with the sequence 5′-TUAAAAUCUACAGUCAUAGGAAU-3′, where the 5′-terminal nucleotide is a 2′-*O*-methoxyethyl vinylphosphonate-modified residue and positions 2, 6, 12, and 14 are 2′-fluoro RNA, with the remainder being 2′-*O*-methyl RNA. The sense strand has the sequence 5′-UCCUAUGACUGUAGAUUUUAA-3′, is primarily 2′-*O*-methyl RNA with 2′-fluoro modifications at positions 7, 9, 10, and 11, and features the C7-amino linker at the 3′-end. Both strands contain phosphorothioate linkages at the last two internucleotide positions at both the 3′ and 5′ termini. All modifications and final product integrity were confirmed by LC-MS analysis.

Following purification and characterization, BCN functionalization of the sense strand was performed as follows: 1 equivalent of the 3′-amino-modified *Hprt* sense strand was dissolved in 1 mL of 100 mM sodium tetraborate buffer (pH 8.5). To this solution, 5 equivalents of (1R,8S,9s)-Bicyclo[6.1.0]non-4-yn-9-ylmethyl N-succinimidyl carbonate (BCN-NHS carbonate), dissolved in 0.5 mL DMSO, were added. The reaction mixture was stirred at room temperature until completion, as confirmed by LC-MS analysis. The resulting BCN-functionalized oligonucleotide was purified by strong anion exchange (SAX) chromatography and desalted by reverse-phase HPLC as previously described. The purified BCN-functionalized sense strand was then duplexed with the complementary antisense strand by mixing equimolar amounts of each strand.

### Generation and purification of Clone F-motavizumab

Clone F, formatted as a single-chain variable fragment (scFv), was genetically fused to the C-terminus of the Fc region of motavizumab, a human IgG1 antibody that targets respiratory syncytial virus (RSV) but does not engage any tissue-relevant targets in this study. This bispecific antibody format was described previously (14). The resulting bispecific antibody fused with Clone F had a mutation introduced at its Fc to reduce its effector function (Leu234Ala, Leu235Ala; LALA) and knob-into-hole mutation to generate the asymmetric bispecific antibody (16,17). Motavizumab without Clone F fusion was used as a monoclonal antibody (mAb) shown in the imaging experiment. All antibodies were expressed in Chinese hamster ovary (CHO) cells using the ExpiCHO™ expression system (Thermo Fisher Scientific) in a transient expression format, following the manufacturer’s protocol.

The purification process was described in detail elsewhere (14). Briefly, culture media was loaded with the appropriate column, and the antibody was obtained using a series of washing, elution, and neutralization steps. Purified samples were dialyzed with a formulation buffer and concentrated using Amicon Ultra-15 Centrifugal Filter Units 30K MWCO (EMD Millipore). The purity, host cell protein levels, and endotoxin content of prepared samples were thoroughly monitored and measured prior to *in vivo* experiments.

### Antibody-siRNA conjugation via glycol engineering and SPAAC chemistry

A schematic of *Hprt* siRNA conjugation to the motavizumab × Clone F scFv bispecific fusion protein using a glyco-engineering method is shown in Figure 1B. Conjugation was performed with the GlyCLICK Azide Activation Kit (Genovis, catalog #L1-AZ1-125). Briefly, 132 mg of Clone F fusion (10.45 mg/mL in 20 mM Histidine-HCl, pH 6.0, with 8% sucrose) was buffer-exchanged to Tris-buffered saline (TBS, pH 7.4) using a protein concentrator (50 kDa MWCO) and adjusted to 22 mg/mL (6 mL). Deglycosylation was performed using a GlycINATOR Maxispin column at 37 °C for 2 h. The deglycosylated protein was eluted, concentrated, and confirmed by LC-MS analysis (Figure S1). For azide activation, GalT enzyme, UDP-GalNAz, and MnCl_2_ were added to the deglycosylated Clone F fusion solution and incubated at 30 °C for 24 h, protected from light. The resulting GalNAz-functionalized Clone F fusion was purified and concentrated using a protein concentrator (50 kDa MWCO) and subsequently confirmed by LC-MS analysis to have two GalNAz groups added (Figure S1). Strain-promoted azide–alkyne cycloaddition (SPAAC) conjugation was then performed by adding the BCN-functionalized *Hprt* duplex (as shown in Figure 1B) to the azide-activated Clone F fusion solution. The BCN-*Hprt* duplex was dissolved in TBS, pH 7.4, and the reaction mixture was incubated at 20 °C for 2 h, then stored at 4 °C overnight. The Clone F-*Hprt* conjugates were purified by size exclusion chromatography using a HiPrep 26/60 Sephacryl S-200 HR column. Purified fractions were concentrated and quantified by NanoDrop absorption at 260 nm. The final product consisted of a mixture of Clone F-*Hprt* conjugates, containing either one (DAR1) or two (DAR2) siRNA molecules per Clone F fusion, in an approximate 1:1 ratio in PBS, with an overall recovery of 55% (Figure S2).

**Figure 1.**
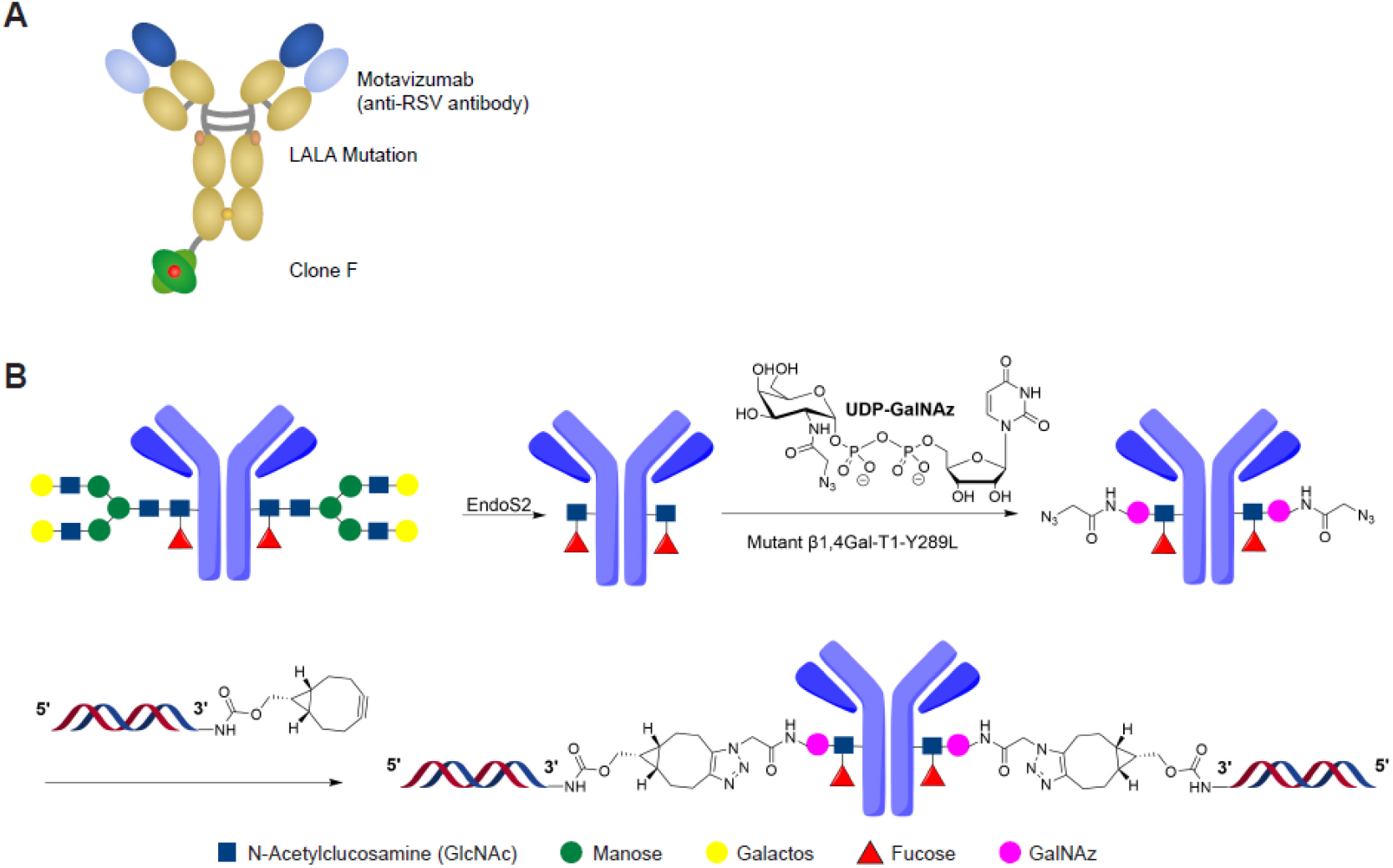
Structure of Clone F-based shuttle and siRNA conjugation method. A) Schematic figure of the Clone F-based shuttle composed of anti-RSV antibody (motavizumab), and Clone F conjugated at the C-terminus of the motavizumab via an amino acid linker. B) Schematic representation of *Hprt* siRNA conjugation to the motavizumab × Clone F fusion using a glycol-engineering approach. The glycan structure depicted is representative of the predominant N-glycans found on motavizumab, which are primarily fucosylated diantennary glycans with variable galactose content.

### ELISA-based binding analysis of Clone F-*Hprt* to IGF1R

To estimate the affinity of Clone F-*Hprt* via enzyme-linked immunosorbent assay (ELISA) the recombinant ectodomains of human and mouse IGF1R were immobilized by adding 100 µL of 4 µg/mL ectodomain in PBS to individual wells of a 96-well maxisorp plate (Thermo Scientific, Cat #243656) and incubated overnight at 4 °C. The following day the plates were washed once with PBS and blocked with the addition of 200 µL of 1% casein (Thermo Scientific, Cat #37528) followed by 1 h incubation. Plates were then washed once with PBS followed by addition of 100 µL per well of a 12-point 1:3 serial dilution (prepared in PBS) of Clone F-*Hprt* from 1000 nM to 0.0056 nM in duplicate wells. Ligands were incubated for 2 h followed by 6 washes in PBS/0.1% tween-20. Plates were tapped dry, then a 1:13000 dilution of HRP-labeled goat anti-human detection antibody (Jackson ImmunoResearch Laboratories, Inc, Cat#109-035-003) was applied to each well. Plates were incubated in detection antibody for 1 h followed by 6 additional washes in PBS/0.1% tween-20. For ELISA development, 100 µL of TMB substrate (Thermo Scientific, Cat #34028) was added to each well and allowed to develop until the highest concentration wells turned moderate-deep blue followed by addition of 100 µL 2N H_2_SO_4_ to stop the reaction. Plates were read on Tecan Spark instrument for absorbance at 450 nm and the resulting values were fitted to a 4-parameter variable slope dose-response model using Prism software to derive EC_50_ values.

### Animal studies of Clone F-*Hprt* by ICV, IV, or SC administration

Adult male C57BL/6NTac mice (Taconic Biosciences, La Jolla, CA) were used for *in vivo* studies. Clone F-*Hprt* solutions were prepared in phosphate-buffered saline and administered via ICV, IV, or SC injection. Animal dosing, sample collection, and analysis described in this manuscript using Clone F-*Hprt* were performed at Ionis Pharmaceuticals under protocols approved by the Institutional Animal Care and Use Committee (IACUC, protocol 2021-1176) at Ionis Pharmaceuticals.

To evaluate the activity of Clone F-*Hprt* in the mouse brain following peripheral administration, mice were dosed IV or SC with Clone F-*Hprt* at 0.3, 1, 3, or 10 mg/kg (siRNA equivalent) on days 1 and 8, at 10 µL/g. As a negative control, unconjugated *Hprt* siRNA was administered IV at the same dose levels and time points. For the positive control, unconjugated *Hprt* siRNA was delivered via ICV injection at 100 µg (in 10 µL) on day 1. ICV injection was done as previously described (18). Mice were sacrificed on day 15, and tissues were collected following terminal euthanasia.

To assess activity following central administration, mice were dosed ICV with either unconjugated *Hprt* siRNA or *Hprt* siRNA conjugated with C16 at the internal P6 position on the sense strand at 10, 30, 100, and 300 μg on day 1, and Clone F-*Hprt* at 3, 10, 30, and 60 μg on day 1. Mice were sacrificed on day 15 and tissues were collected following terminal euthanasia.

### *Hprt* mRNA quantification in tissues by qRT-PCR

Each sample was homogenized in 1 mL of Trizol reagent (Thermofisher scientific, Waltham, MA) and total RNA was extracted using a mini-RNA purification kit (Qiagen, Valencia, CA), according to manufacturer’s protocol. The Life Technologies ABI QuantStudio 7 Flex Sequence Detection System (Applied Biosystems Inc, Carlsbad CA) was employed for realtime quantitative PCR analysis (rt-qPCR). Briefly, 10 µL RT-PCR reactions containing 400 ng of RNA were run with the AgPath-ID One-Step qRT-PCR Kit (Thermofisher scientific, Waltham, MA) reagents with primer probe sets listed below mouse *Hprt* and *Ppia*, a ubiquitously expressed housekeeping gene. All qPCR reactions were run in technical triplicates; outliers in triplicates, which resulted in > 20% coefficient variation of sample average, were excluded. The expression level of total *Hprt* mRNA was normalized to that of *Ppia* mRNA, and this was further normalized to the level measured in control animals that were administered PBS. *Hprt* mRNA levels are reported as percent control.

### Primer probe sets used for qPCR

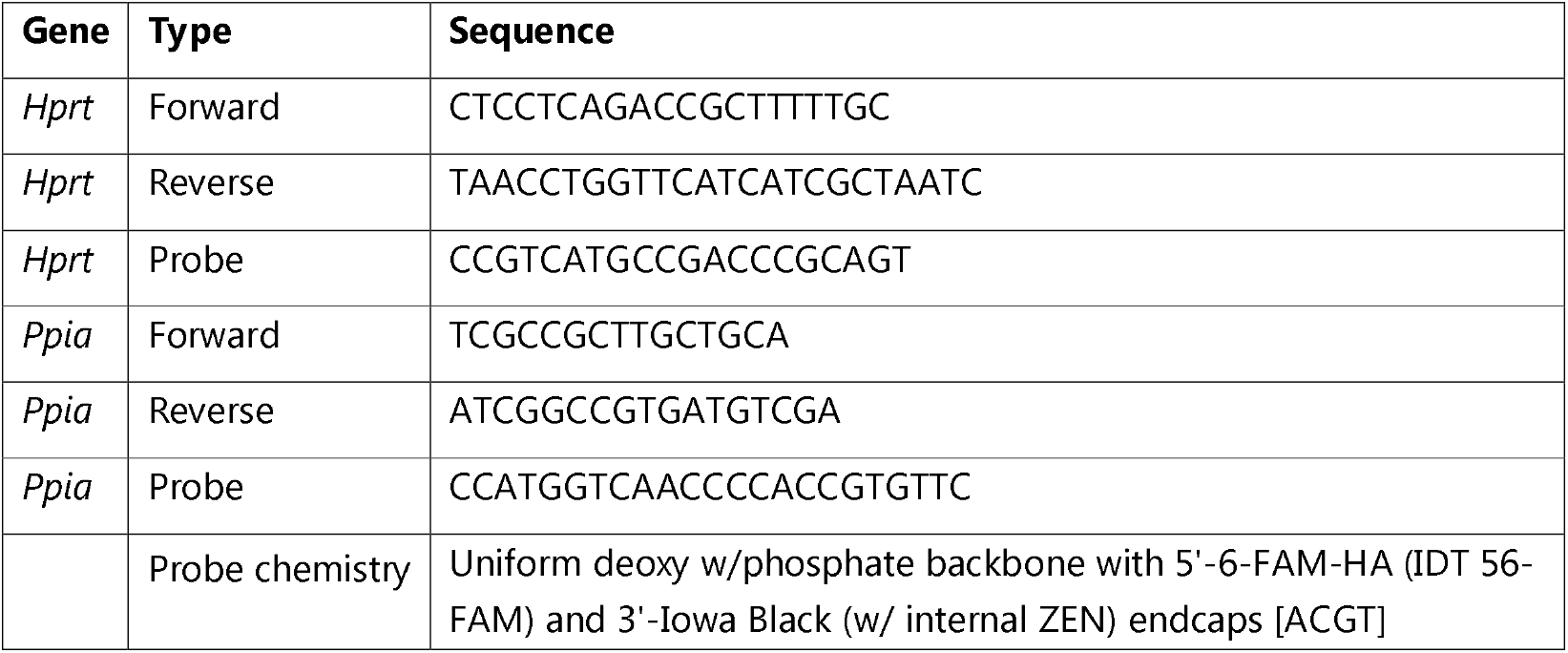

#### LC-MS quantification of *Hprt* siRNA in mouse liver, kidney, and lung

All tissue samples were extracted using a liquid-liquid extraction (LLE) method, followed by solid-phase extraction (SPE) and analysis using liquid chromatography-mass spectrometry (LC-MS). Naïve liver homogenate was used for a calibration curve and QCs to quantitate liver and kidney. Naïve lung homogenate was used for a calibration curve to quantitate lung.

Approximately 50 mg tissue sample was cut and weighed into a well in a 96 deep well (2 mL) plate containing approximately 0.25 cm^3^ Matrix Green homogenization beads. Homogenization buffer (20 mM Tris pH 8, 20 mM EDTA, 0.1 M NaCl, 0.5% NP40) was added to a final volume of 500 µL to each well, and the samples were homogenized using the Mini Bead Beater (Thomas Scientific, Swedesboro, NJ). Samples were subsequently treated with ammonium hydroxide (300 µL, J.T. Baker/VWR 9721-02) and extracted by LLE, adding 800 µL phenol:chloroform:isoamyl alcohol (25:24:1)[Sigma Aldrich], followed by a SPE using a 96-well Strata X packed plate (Phenomonex Inc., CA) and a final pass through using a Protein Precipitation Plate (Phenomonex Inc., CA). Eluates were dried down under nitrogen at 50 °C and reconstituted with water containing 100 µM EDTA.

The reconstituted samples were analyzed by ion-pairing (IP) LC-MS/MS on an Agilent 6495 triple quadrupole system (Agilent, Wilmington, DE, USA). Samples were injected and separated on an ACQUITY UPLC Oligonucleotide BEH C18 column ((130 A, 1.7 µm, 2.1×50 mm) Waters, Milford, MA, USA) heated to 70 °C, with a flow rate of 0.3 mL/minute. A binary solvent system with mobile phase A (0.5% (47.6 mM) 1,1,1,3,3,3-Hexafluoro-2-propanol (HFIP)/ and 0.17% (13.15 mM) *N,N*-diisopropylethylamine (DIPEA); mobile phase B (Methanol (MeOH)) was used. A gradient from 15 to 40% MeOH over 6 min was used to elute analyte.

All mass measurements were made on-line. The antisense strand calculated concentrations were used as a representative of the concentration of dosed conjugated siRNA in tissue. Mass spectra were obtained using a nozzle voltage of −1500 V, a nebulizer gas flow of 15 psi, a sheath gas flow rate of 10 L/min at 350 °C, a drying gas flow rate of 5 L/min at 350 °C, and a capillary voltage of −4000 V. Chromatograms were analyzed using the Agilent Mass Hunter software. Concentrations of analytes were quantitated and calculated as antisense strand levels, expressed as unconjugated duplex equivalents, using the corresponding calibration curve with a quantitation range of 0.03 µg/mL (0.002 µM) to 1473 µg/mL (100 µM).

#### HELISA quantification of *Hprt* siRNA in mouse brain, spinal cord, and quadriceps

Briefly, samples were weighed into individual wells of a 2 mL 96-well plate; then 500 µL homogenization buffer (20 mM Tris pH 8, 20 mM EDTA, 0.1 M NaCl, 0.5% NP40) was added to those corresponding wells. Control tissue homogenate for curves and QCs was made by weighing control mouse brain (or control mouse liver) and adding homogenization buffer at a 25 to 1 ratio, so that 500 µL of homogenate contained 20 mgs of tissue. Approximately 0.25 cm^3^ granite beads were added to the wells containing samples and calibration standards. 500 µL aliquots were pipetted into a 96-well plate and appropriate amounts of calibration standards were spiked in the corresponding wells. The curve and QCs were spiked with unconjugated *Hprt*. The plates were then extracted via a liquid-liquid extraction with ammonium hydroxide and phenol:chloroform: isoamyl alcohol (25:24:1). The aqueous was then dried down under vacuum.

Tissue, curve, and QC samples were reconstituted using “reconstitution buffer” (0.1% K_2_EDTA, 0.5% Tween 20) and further diluted in reconstitution buffer, as necessary. 25 µL of sample and 25 µL of control human plasma (for running matrix) were aliquoted into wells and 450 uL of 1.875 nM probe solution was added. The hybridization ELISA method was carried out and read on a Molecular Devices Spectramax Gemini XPS (Sunnyvale, CA). The assay involves binding of the analyte antisense strand to the sense DNA probe and subsequent detection of the double-strand hybridized drug using an antibody that recognizes a 5’ (or 3’)-linked digoxigenin. Selection of the double-strand hybrid is accomplished by elimination of single-strand (un-hybridized probe) using S1 nuclease digestion. Sample quantitation was calculated from the co-extracted calibration curve using SoftMax Pro (Molecular Devices, LLC). The range of the assay for the antisense strand (which represents the duplex siRNA), was ∼0.00375-0.5 μM (∼0.055 - 7.364 µg/g of unconjugated duplex) in 20 mg of control monkey brain (for all cortex and spinal cord sections) or liver (for quadriceps). Curve and QC samples quantified within 20% of their nominal values. The %CV between duplicate wells on the plate was <20% for all curves, QC’s and samples. All samples were stored at −20 °C ± 5 °C, upon receipt.

#### Mouse brain imaging of *Hprt* siRNA and HPRT protein

##### Tissue processing for immunohistochemistry

Tissues were fixed in 10% formalin for 48-72 h, embedded, and sectioned at 4 µm.

#### Immunohistochemical detection of HPRT protein

Slides were stained with rabbit monoclonal HPRT antibody (ThermoFisher, PA5-22283) on a Ventana Ultra staining system. Slides were treated with heat-induced antigen retrieval (HIER) with Ventana CC1 solution (Ventana, 950-500) for 64 min. Tissues were permeabilized with 5% Tween 20 (VWR, 0777-1L) for 32 min at 37 °C. The primary antibody was diluted with Discovery Antibody Diluent (Ventana, 760-108) at 1:25 for heart and muscle and 1:800 for all other tissue types, then incubated for 12 h at room temperature. The antibody was detected with UltraMap anti-rabbit polymer detection (Ventana, 760-4315) and visualized with the Ventana ChromoMap DAB kit (Ventana, 760-159). Images were scanned on a Hamamatsu S360 scanner at 20× resolution.

#### Immunofluorescence and confocal imaging of human IgG in mouse brain

Drug treatment and tissue collection were conducted at NDIC. Eight-week-old C57BL/6J mice were treated with either motavizumab at a 30 mg/kg dose or Clone F-fused motavizumab at its molar equivalent dose (35.1 mg/kg) via a single IV administration. After 48 h, the mice were deeply anaesthetized using isoflurane, followed by transcardial perfusion to avoid residual blood in brains. Brains were collected and fixed immediately with 4% paraformaldehyde, followed by embedding into a 30% sucrose solution.

Each brain was embedded in Optimal Cutting Temperature (OCT) compound and rapidly frozen. Frozen brains were sectioned into 40-μm slices using a cryostat microtome. For antigen retrieval, sections were treated with HistoVT One (Nacalai Tesque, 06380-05) according to the manufacturer’s instruction, followed by permeabilization with 0.2% Triton X-100 for 30 min at room temperature. Autofluorescence was quenched using TrueBlack Lipofuscin Autofluorescence Quencher for 30 sec. Sections were then blocked with 5% normal goat serum for 1 h at room temperature. Primary staining was performed with anti-human IgG antibody (SouthernBiotech, 6145-01; 1:300 in blocking solution) overnight at 4 °C. The following day, sections were washed with PBS and incubated with Alexa Fluor 594 anti-rabbit IgG secondary antibody (Invitrogen, A32740; 1:500) for 2 h at room temperature. After staining, sections were mounted on glass slides using mounting medium (Vector Laboratories, H-2000) and imaged with a confocal microscope (LSM900, Zeiss, Germany). Z-stack images were acquired at 1-μm intervals across 10 optical sections using a 20× objective with a 3×3 tile scan.

#### Western blot detection of IGF1R in tissue lysates

Four adult female C57BL/6NTac mice (Taconic Biosciences, La Jolla, CA) were euthanized under deep isoflurane anesthesia. Brain, spinal cord, and peripheral organs, including muscle, liver, lung, heart, kidney, and spleen, were collected, snap-frozen in liquid nitrogen, and stored at −80 °C. Dissected tissues were weighed and homogenized using a bullet blender with tungsten beads in radioimmunoprecipitation assay (RIPA) buffer supplemented with protease and phosphatase inhibitors (Roche) at 4 °C. Samples were then sonicated (30% amplitude, 1-second pulse on/off cycle for 1 minute) and clarified by centrifugation at 12,000-14,000 rpm for 10 min.

A total of 20μg of protein per well was separated on 4-20% gradient SDS-polyacrylamide gels and transferred to nitrocellulose membranes. Membranes were blocked with 5% non-fat milk in Tris-buffered saline (TBS) and probed with an anti-IGF1R β-chain antibody (Abcam, Cat# ab182408; 1:1,000 dilution). Lysates from IGF1R knockout HeLa cells (IGF1R Knockout Cell Line-HeLa #4, Creative Biogene, Order# CBGNW0010) were included as a negative control. These cells were cultured in DMEM (Gibco, Cat# 10566-016) supplemented with 10% FBS (Gibco, Cat# 10099-141) and 1× Antibiotic-Antimycotic (Gibco, Cat# 15240-062).

Initially, GAPDH was used as a loading control (anti-GAPDH antibody, Cell Signaling Technology, Cat# 5174S; 1:1,000), but due to high variability in band intensity across tissues, total protein staining using No-Stain™ Protein Labeling Reagent (Invitrogen, Cat# A44449) was used instead. Protein bands were visualized using the ChemiDoc MP Imaging System (Bio-Rad) and quantified with Image Lab software (Bio-Rad, version 6.1.0, build 7).

### Statistical analysis

Statistical significance between groups was assessed using one-way analysis of variance (ANOVA) followed by Dunnett’s multiple comparisons test to compare treatment groups against the control. Significance was defined as p < 0.05.

## RESULTS

### Generation of Clone F-*Hprt* by conjugating the bispecific antibody with siRNA

We conjugated the motavizumab × Clone F bispecific fusion antibody to an siRNA targeting hypoxanthine-guanine phosphoribosyl transferase (*Hprt*), a housekeeping gene. The bispecific antibody was generated by fusing an anti-IGF1R single-chain variable fragment (scFv) to the C-terminus of motavizumab, a human IgG1 antibody against respiratory syncytial virus (RSV) used here without tissue-relevant binding (14), (19), as described in Materials and Methods (Figure 1A). To minimize undesired Fc-mediated effector functions, an Fc mutation (Leu234Ala, Leu235Ala; LALA) was introduced (17).

The *Hprt*-targeting siRNA was synthesized, functionalized, and conjugated to the Fc region of the bispecific fusion antibody using a glyco-engineering approach, as described in the Materials and Methods. Briefly, the process involved antibody deglycosylation followed by site-specific azide activation to enable covalent attachment of the siRNA (Figure 1B).

### Clone F-*Hprt* retains binding to IGF1R after conjugation

We evaluated the binding of Clone F-*Hprt* to IGF1R to ensure its binding to RMT target was preserved after the conjugation. A sandwich ELISA with human or mouse IGF1R ectodomains clearly demonstrated sigmoidal binding of the conjugates with nanomolar EC_50_ values (EC_50_: human IGF1R - 15 nM; mouse IGF1R - 119 nM; Figure S3). These data confirm that the final conjugate, Clone F-*Hprt*, retained its binding to IGF1R after the siRNA conjugation.

### Clone F-*Hprt* retains activity and reduces *Hprt* expression after ICV administration

To assess whether Clone F conjugation enhances the activity of *Hprt* siRNA in the brain following central administration, Clone F-*Hprt* was delivered via ICV injection to 8-week-old C57BL/6NTac mice. The study included a control group receiving PBS, along with groups treated with either unconjugated *Hprt* siRNA or a lipophilic C16-conjugated *Hprt* siRNA with the C16 installed at the internal P6 position on the sense strand (C16-*Hprt*). C16-siRNAs have shown broad CNS distribution and uptake after CSF delivery, making C16-*Hprt* a relevant benchmark for the ICV dosing context (6). The unconjugated and C16-conjugated *Hprt* siRNA groups received doses of 10, 30, 100, or 300 μg, whereas Clone F-*Hprt* was evaluated at 3, 10, 30, or 60 μg due to compound availability and the practical constraints of a large protein conjugate.

During the two weeks post-treatment, all animals exhibited normal home cage behavior, and their weight gain was comparable to the PBS groups (data not shown). On day 14, tissues were collected from multiple CNS regions, and RNA was extracted to quantify *Hprt* mRNA levels. The unconjugated *Hprt* siRNA showed a dose-dependent reduction in *Hprt* mRNA, and this effect was enhanced by both C16-*Hprt* and Clone F-*Hprt* (Figure 2A). We calculated ED_50_ values for each compound and compared the fold increase in potency relative to unconjugated *Hprt* siRNA (Figure 2B,C and Figure S4). As expected, C16-*Hprt* substantially improved potency across all CNS regions. For example, in the cortex, the ED_50_ of C16-*Hprt* was 19.2 μg (95% CI: 7.9-33.2), whereas the ED_50_ of unconjugated *Hprt* siRNA was 117.9 μg (92.4-154.0), corresponding to a 6.1-fold improvement. Fold improvements with C16-*Hprt* ranged from 2.3-fold in the olfactory bulb to 33.8-fold in the striatum (Figure 2C and Figure S4). Clone F-*Hprt* also enhanced siRNA potency, but to a lesser extent than C16-*Hprt*. In the cortex, the ED_50_ of Clone F-*Hprt* was 28.3 μg (19.0-46.8), a 4.2-fold improvement compared to unconjugated siRNA. Improvements with Clone F-*Hprt* were observed in multiple regions, including 3.8-fold in the hippocampus, 3.0-fold in the striatum, 3.3-fold in the thalamus, and 2.4-fold in the cerebellum (Figure 2C and Figure S4). In contrast, Clone F-*Hprt* did not improve potency in the brainstem (0.9-fold) or spinal cord (0.8-fold), likely reflecting limited diffusion due to its molecular size or chemical properties (Figure S4).

**Figure 2.**
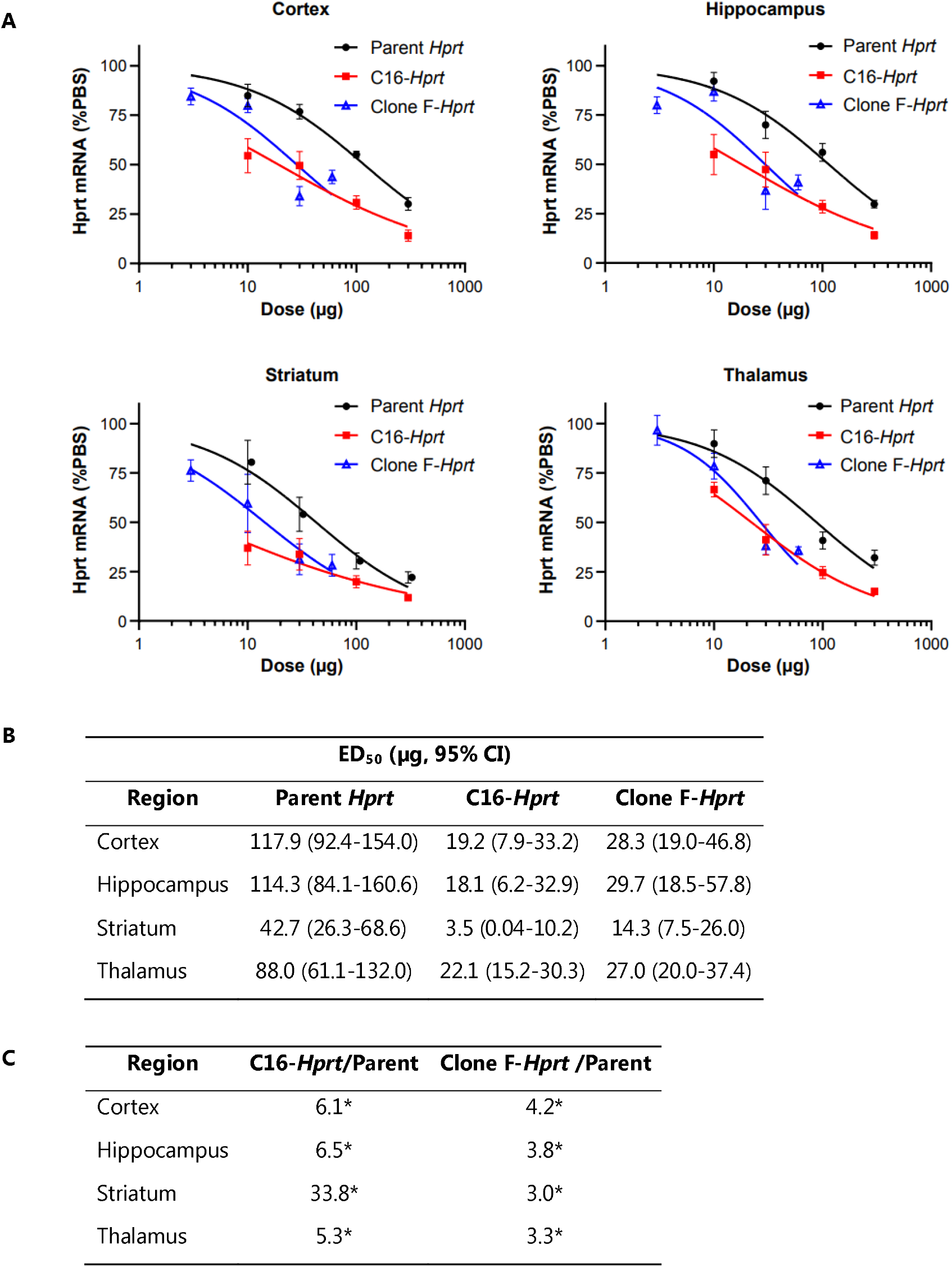
Dose-dependent reduction of *Hprt* mRNA across multiple CNS regions following ICV administration of Clone F-*Hprt*, C16-*Hprt*, or unconjugated *Hprt* siRNA. (A) Dose-response curves for *Hprt* mRNA levels after ICV administration at the indicated doses. Levels were quantified by qRT-PCR from tissues collected two weeks post-dosing. Curves were generated using non-linear regression (log[agonist] vs. response) in GraphPad Prism, with the bottom constraint set to 0 and the top to 100. Data represent mean ± SEM. The unconjugated *Hprt* siRNA and C16-*Hprt* groups consisted of six animals per group (n = 6), while the Clone F-*Hprt* group consisted of four animals per group (n = 4). (B) ED_50_ values (µg, 95% CI) for unconjugated *Hprt* siRNA, C16-*Hprt*, and Clone F-*Hprt* in cortex, hippocampus, striatum, and thalamus. (C) Fold improvements in potency of C16-*Hprt* and Clone F-*Hprt* relative to unconjugated *Hprt* siRNA across CNS regions. Asterisks denote a statistically significant difference (p < 0.05) in the potency (ED_50_) of the conjugate compared to the unconjugated parent siRNA, as determined by statistical comparison of the dose-response curves.

### Clone F-*Hprt* mediates CNS knockdown following IV and SC delivery

We then investigated whether Clone F-*Hprt* could mediate gene knockdown in the CNS following peripheral administration. To test this, PBS, unconjugated *Hprt* siRNA, and Clone F-*Hprt* were administered IV or SC at doses of 0.3, 1, 3, and 10 mg/kg on days 1 and 8. CNS tissues were collected on day 15. A single ICV dose of 100 μg unconjugated *Hprt* siRNA was included as a positive control, corresponding to the previously determined ED_50_ of 117 μg in the cortex. All animals maintained normal home cage behavior and exhibited weight gain comparable to controls during the two-week post-dosing period (data not shown). Bulk RNA analysis revealed that ICV administration of unconjugated *Hprt* siRNA (100 μg) reduced *Hprt* mRNA levels to 25-50% of baseline across CNS regions, corresponding to a 50-75% knockdown, as confirmed by qRT-PCR (Figure 3A). However, IV administration of unconjugated siRNA (10 mg/kg) did not produce any knockdown (Figure 3A; Figure S5A). Notably, IV administration of Clone F-*Hprt* (0.3-10 mg/kg) produced a clear dose-dependent reduction in *Hprt* mRNA across CNS regions, including deep brain structures such as the striatum and thalamus (Figures 3A and 3B). In additional regions, such as the brainstem and spinal cord, significant knockdown was observed at higher doses, whereas the olfactory bulb showed only a non-significant trend (Figures S5B and S5C). While SC administration of Clone F-*Hprt* was less effective, it still produced modest *Hprt* knockdown compared with IV dosing at the highest dose (10 mg/kg) (Figure 3C; Figure S6).

**Figure 3.**
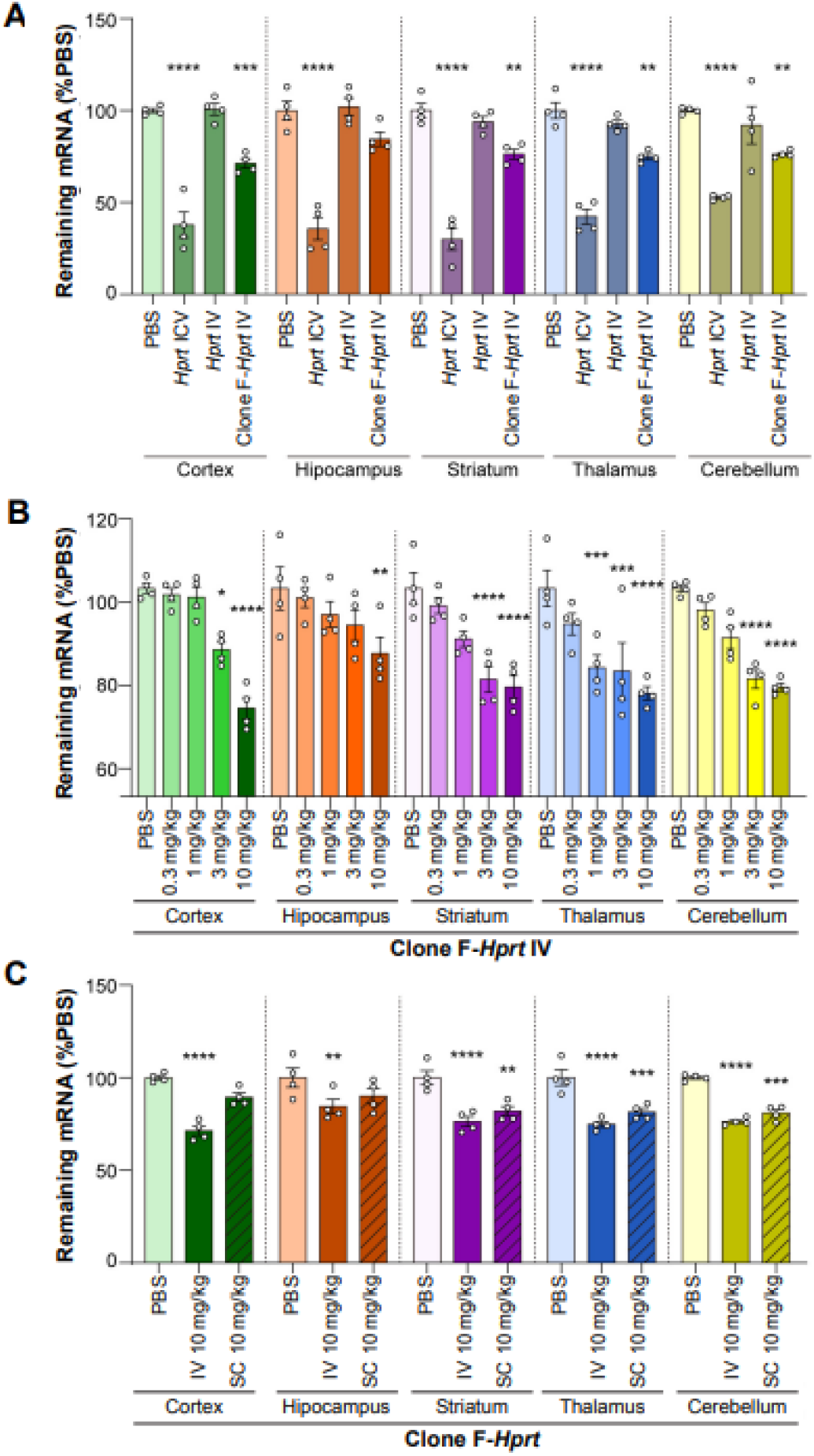
mRNA reduction across multiple CNS regions after IV administration of unconjugated *Hprt* siRNA, IV or SC administration of Clone F-*Hprt*. Mice received PBS, unconjugated *Hprt* siRNA, or Clone F-*Hprt* by IV or SC injection (0.3-10 mg/kg, days 1 and 8); tissues were collected on day 15. A single ICV dose of 100 μg siRNA served as a positive control. *Hprt* mRNA levels were measured by qRT-PCR and expressed relative to PBS controls (mean ± SEM; n = 4). (A) Remaining *Hprt* mRNA after ICV siRNA (100 μg), IV siRNA (10 mg/kg), or IV Clone F-*Hprt* (10 mg/kg). (B) Dose-dependent knockdown after IV Clone F-*Hprt*. (C) Comparison of IV vs SC Clone F-*Hprt* at 10 mg/kg. Significance was determined by one-way ANOVA with Dunnett’s multiple comparisons test (*p < 0.05; **p < 0.01; ***p < 0.005; ****p < 0.001).

To confirm that the enhanced gene knockdown was due to improved brain delivery with the Clone F-based shuttle, we designed a separate study to evaluate antibody distribution in mouse brain tissues. Motavizumab (30 mg/kg) and Clone F-fused motavizumab (35.1 mg/kg) were administered IV to mice at equimolar ratios, as described in the Methods. Brain tissues were collected 48 h post-dose because human immunoglobulin (hIgG) signal in the brain declines rapidly and becomes difficult to detect by one week following IV or SC administration. Immunofluorescence and confocal imaging of hIgG in brain sections clearly demonstrated that Clone F-fused motavizumab achieved markedly greater localization than motavizumab, confirming delivery to both brain microvessels and parenchymal cells. At this 48-h time point, enhanced hIgG signals from Clone F-fused motavizumab were consistently observed across four brain regions (cortex, hippocampus, hypothalamus, and thalamus; Figure 4).

**Figure 4.**
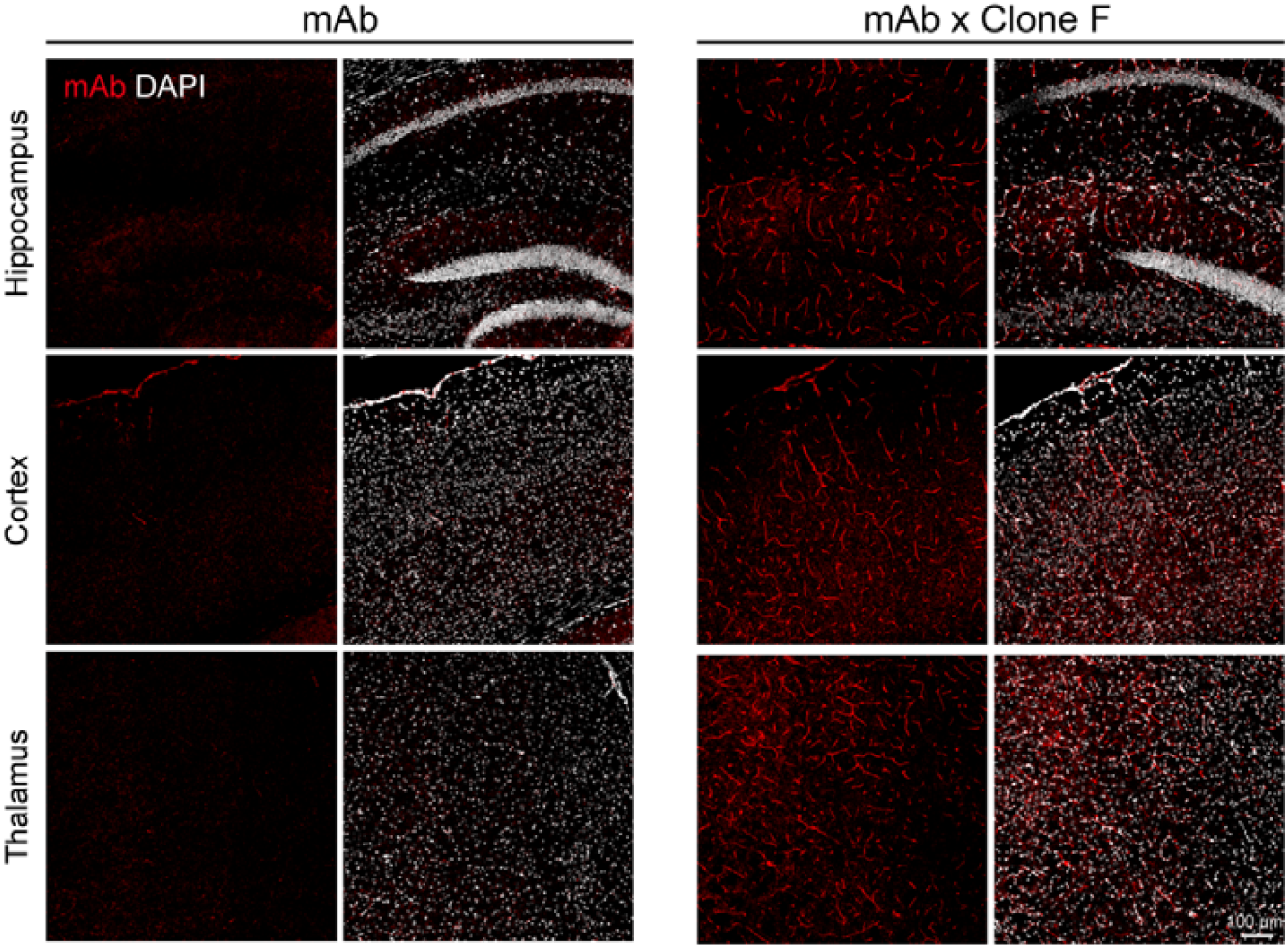
Improved brain distribution of Clone F-fused motavizumab compared to parental motavizumab. Immunofluorescence and confocal imaging of hIgG (red) revealed markedly higher signals for Clone F-fused motavizumab in cortex, hippocampus, hypothalamus, and thalamus, confirming enhanced delivery to both brain microvessels and parenchymal cells. Nuclei were counterstained with DAPI (white). Scale bar, 50 μm.

Next, we sought to determine whether the increased efficacy of Clone F-*Hprt* was attributed to Clone F-mediated delivery of siRNA into CNS tissues. As an example, we examined cortex as a representative CNS region. As referenced previously, unconjugated *Hprt* siRNA was barely detectable in the cortex even after IV dosing at 10 mg/kg. In contrast, Clone F-*Hprt* treatment produced a clear, dose-dependent increase in cortical siRNA levels, as quantified by HELISA (Figure 5A and 5C). Concentrations rose from 0.008 µg/g at 0.3 mg/kg to 0.187 µg/g at 10 mg/kg. For comparison, ICV administration of 100 µg unconjugated siRNA resulted in cortical concentrations of ∼1.388 µg/g, indicating that Clone F-*Hprt* conjugate enabled IV delivery to achieve brain exposures approximately 7-fold lower than direct ICV dosing. Importantly, although these IV exposures are lower than those achieved by ICV, recent clinical trial data demonstrate that sub-µg/g concentrations in CNS tissue can still produce functional knockdown, and in our study, the 0.187 µg/g level at the highest IV dose was associated with a modest but measurable activity in the cortex (∼29% reduction), supporting the relevance of these exposures (Figure 5C). Immunohistochemical staining for HPRT protein (Figure) confirmed target engagement, showing reduced HPRT immunoreactivity in the cortex.

**Figure 5.**
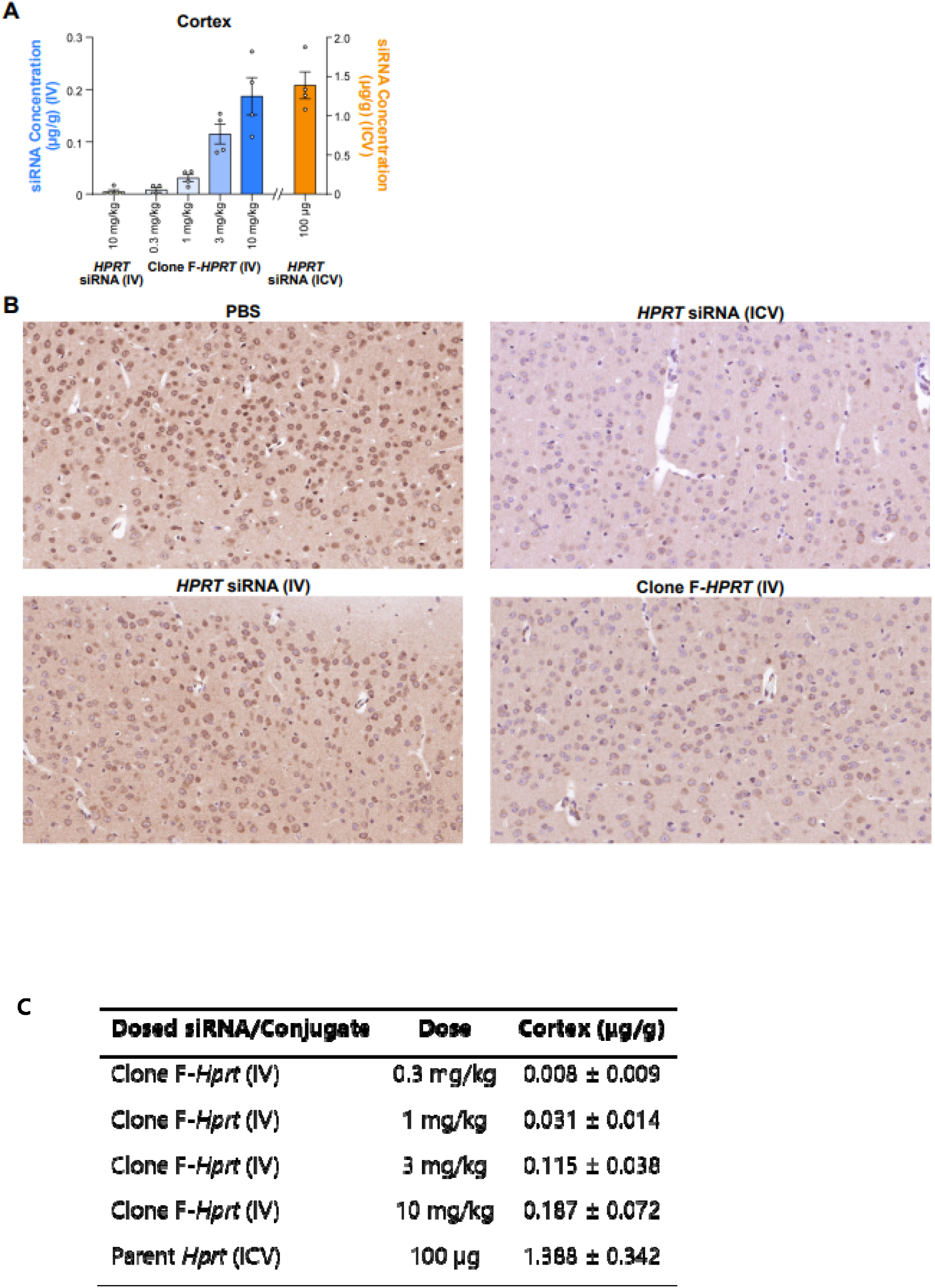
Improved delivery of siRNA into the cortex by Clone F-*Hprt* conjugate. (A) HELISA-based quantification of *Hprt* siRNA in cortex following IV administration of Clone F-*Hprt* at 0.3-10 mg/kg or ICV administration of 100 µg unconjugated siRNA. Data are presented as mean ± SD. (B) Immunohistochemical staining of HPRT protein in cortex showing strong reduction with ICV administration of unconjugated *Hprt* siRNA (100 µg), and evident knockdown with IV Clone F-*Hprt* (10 mg/kg), compared with PBS or IV unconjugated siRNA (10 mg/kg). Scale bar, 100 μm. (C) Tabulated summary of cortical siRNA concentrations (mean ± SD, n = 4) for Clone F-*Hprt* following IV administration (0.3–10 mg/kg) and for unconjugated Hprt siRNA following ICV administration (100 µg).

High doses of oligonucleotide therapeutics, including siRNAs, have been associated with safety concerns in some tissues (5). To evaluate potential CNS effects in our study, we assessed the expression of cluster of differentiation 68 (*Cd*68) and glial fibrillary acidic protein (*Gfap*), marker genes related to inflammatory microglia or macrophages (20) and astrocytes (21), respectively. Gene expression levels in groups treated with either unconjugated *Hprt* siRNA or Clone F-*Hprt* were comparable to those in PBS-treated controls (Figure S7), indicating no evidence of neuroinflammation under the conditions tested.

Taken together with the biodistribution and activity data, these findings demonstrate that anti-IGF1R antibody shuttle enables effective and well-tolerated delivery of *Hprt* siRNA into the CNS with both IV and SC dosing. This supports IGF1R as a validated target for mediating oligonucleotide transport across the BBB.

### Clone F-*Hprt* induces knockdown in peripheral tissues following IV and SC delivery

We next assessed the remaining mRNA levels and siRNA levels in six peripheral tissues (quadriceps, liver, heart, lung, kidney, and spleen) from the same animals, whose brains were analyzed in the previous section. Figure 6A summarizes results from a single 10 mg/kg dose of unconjugated *Hprt* siRNA or Clone F-*Hprt* given IV or SC, as well as from ICV dosing. Following IV administration of 10 mg/kg dose of unconjugated *Hprt* siRNA, only modest knockdown was observed in quadriceps (53.4 ± 3.0% of control) and liver (62.6 ± 7.4%), with little or no activity in heart, lung, kidney, or spleen (Figure 6A). In contrast, the same dosage of Clone F-*Hprt* produced robust reductions in quadriceps (22.0 ± 1.4%), liver (10.9 ± 2.6%), and heart (38.3 ± 1.9%), with more modest effects in lung (62.4 ± 5.6%). SC administration of Clone F-*Hprt* at 10 mg/kg also induced significant knockdown in quadriceps, liver, lung, and heart, although the extent was consistently less pronounced than IV dosing. As expected, ICV administration resulted in minimal knockdown in peripheral tissues, confirming limited systemic distribution via this route (Figure 6A). We next evaluated dose-response relationships for IV and SC administration of Clone F-*Hprt* (Figure 6B). IV dosing yielded ED_50_ values of 0.3 mg/kg in quadriceps, 1.0 mg/kg in liver, and 4.0 mg/kg in heart, whereas SC dosing produced a right-shifted curve with lower overall efficacy, indicating reduced potency and delivery efficiency via the SC route. Notably, quadriceps knockdown reached ∼50% already at the lowest effective IV dose (0.3 mg/kg), underscoring the high sensitivity of muscle tissue to Clone F-*Hprt*. In contrast, Clone F-*Hprt* had little or no activity in kidney and spleen regardless of route, suggesting that siRNA exposure in these tissues did not translate into effective knockdown. (Figure 6A, Figure S8).

**Figure 6.**
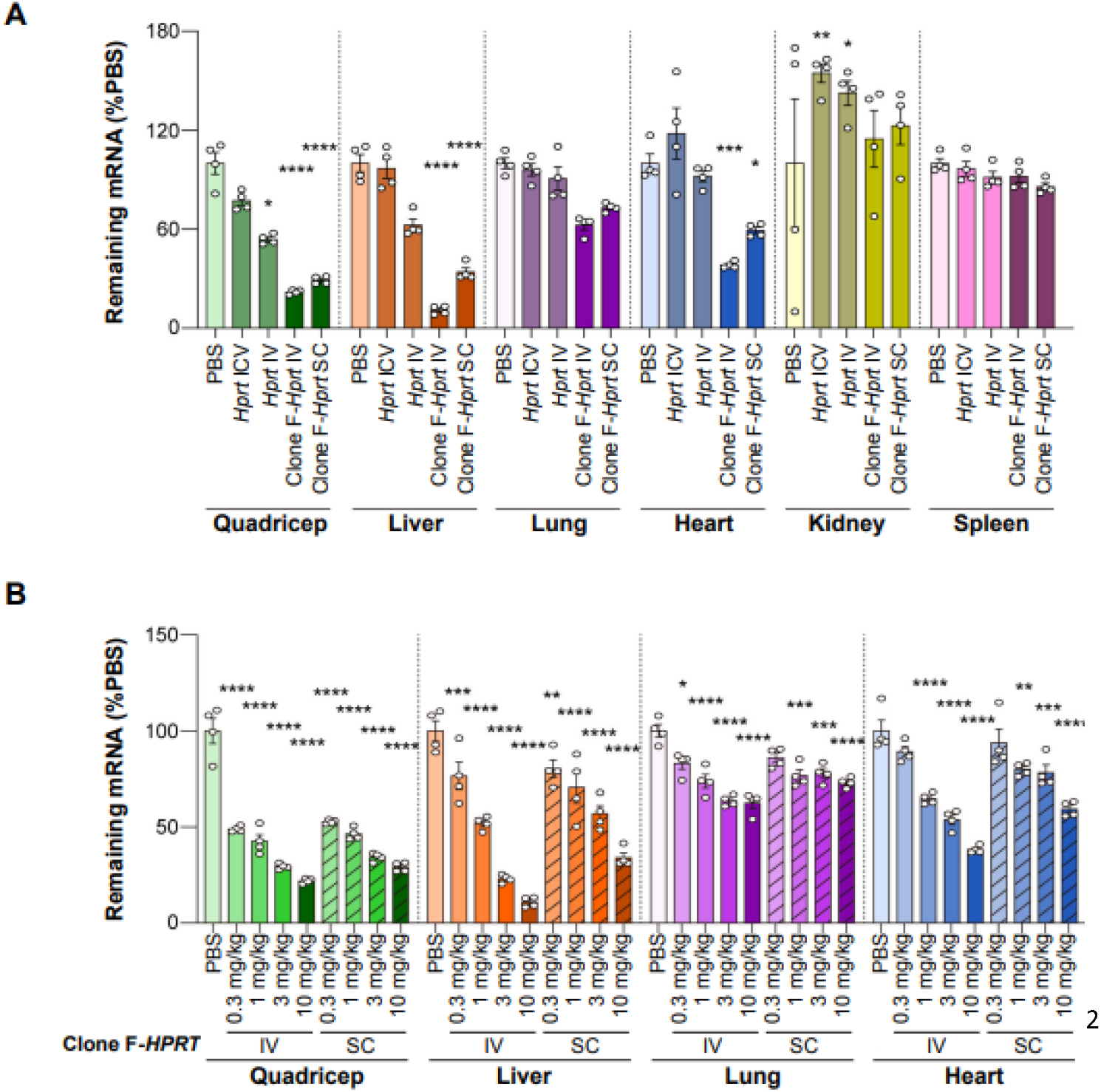
*Hprt* mRNA levels in peripheral tissues after administration of unconjugated siRNA or Clone F-*Hprt*. Peripheral tissues were collected one week after the second weekly dose. (A) Remaining *Hprt* mRNA levels in quadriceps, liver, lung, heart, kidney, and spleen after IV or SC dosing of Clone F-*Hprt* (10 mg/kg) and IV or ICV dosing of unconjugated siRNA (10 mg/kg or 100 µg, respectively) expressed as % of PBS-treated controls (mean ± SEM; n = 4). (B) Dose-dependent reduction in *Hprt* mRNA levels in quadriceps, liver, lung, and heart after IV or SC administrations of Clone F-*Hprt* at doses from 0.3 to 10 mg/kg. Statistical analysis was conducted using ordinary one-way ANOVA with Dunnett’s multiple comparisons test. *: P < 0.05; **: P < 0.01; ***: P < 0.005; ****: P < 0.001.

Given the relatively high expression of IGF1R in the brain (14), the strong potency of Clone F-*Hprt* in various peripheral tissues was unexpected. To investigate this further, we quantified siRNA concentrations in these tissues. Tissue concentrations were measured in quadriceps (by HELISA) and in liver, lung, and kidney (by LC-MS, as described in Materials and Methods). All values are expressed as unconjugated siRNA duplex equivalents (µg/g) for comparability and are presented together with the graphical data in Figure 7. Concentrations in heart were not determined; however, mRNA knockdown and immunohistochemistry were assessed in this tissue (Figures 6 and 7B). These quantitative concentration data provide a framework to directly relate tissue exposure to the degree of mRNA knockdown.

**Figure 7.**
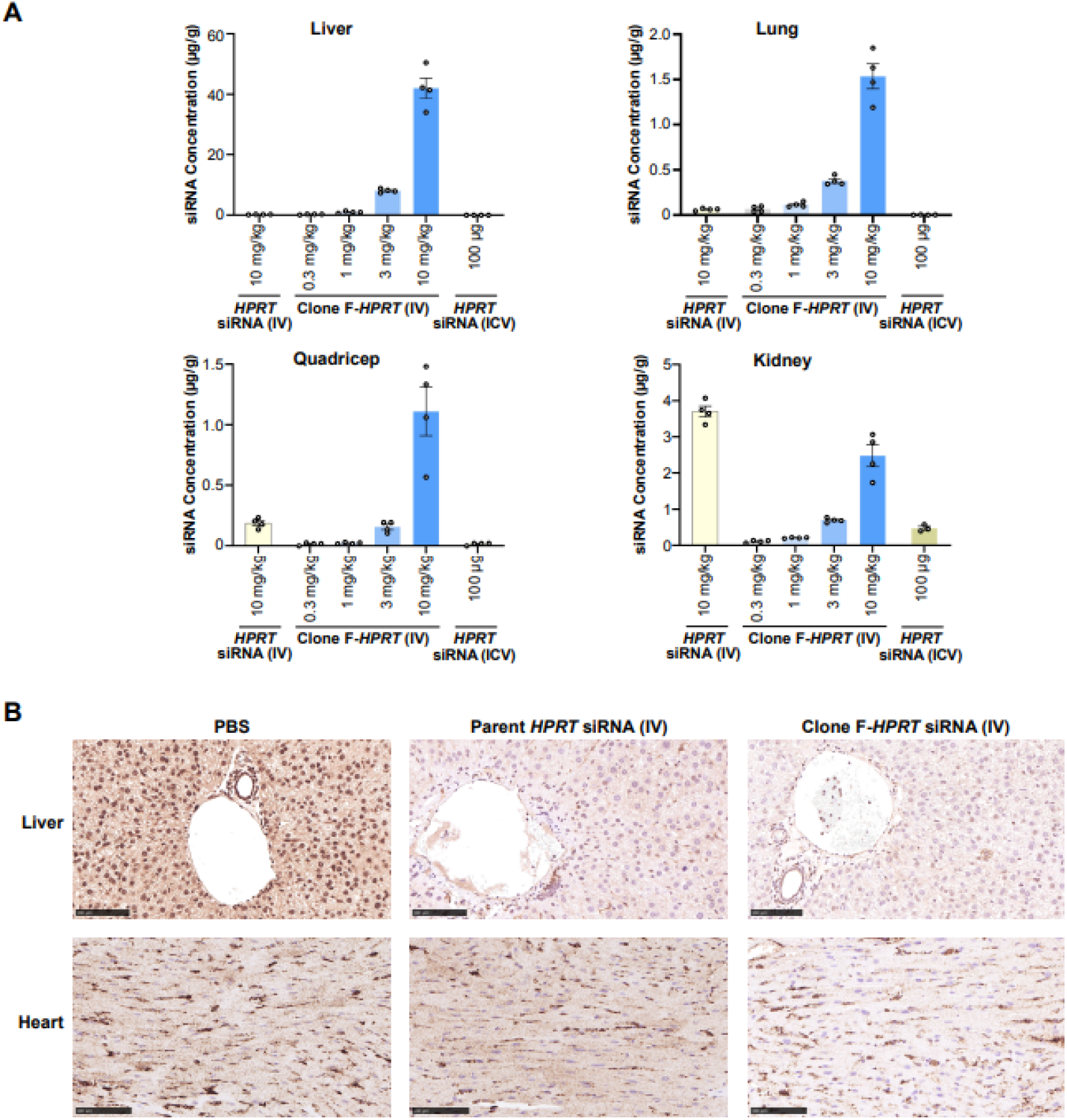
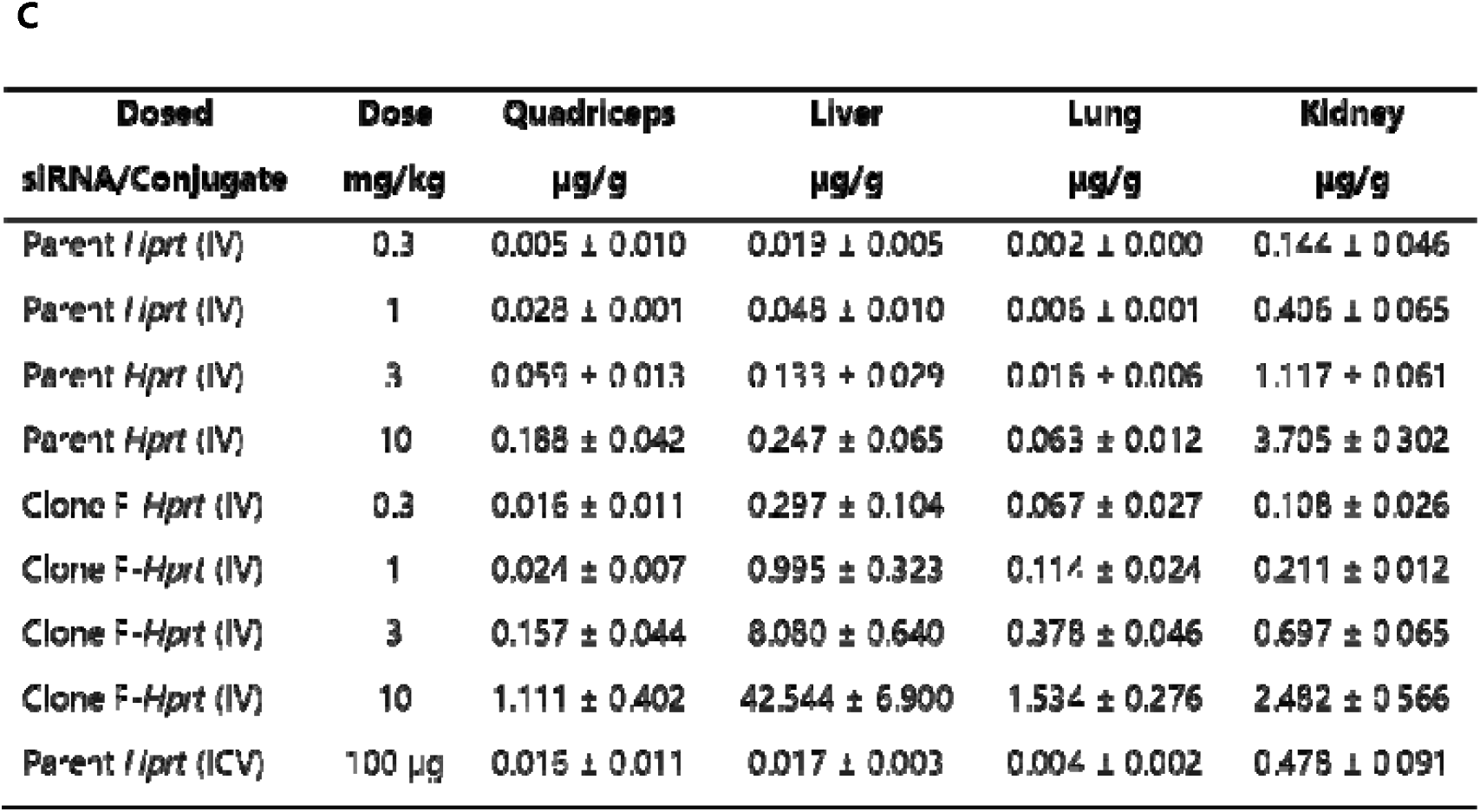
siRNA concentrations and target protein levels in peripheral tissues after IV administration of Clone F-*Hprt. (A) Tissu*e concentrations in liver, lung, quadriceps, and kidney showing dose-dependent accumulation of Clone F-Hprt, with kidney levels reduced compared to unconjugated siRNA. (B) Immunohistochemistry of HPRT protein in liver and heart tissues after IV administration of PBS, unconjugated *Hprt siRNA, or C*lone F-*Hprt. Scale bar*, 100 *μ*m. (C) Summary of tissue concentrations (mean ± SD, n = 4) in quadriceps (measured by HELISA) and in liver, lung, and kidney (measured by LC-MS, as described in Materials and Methods). All values are expressed as unconjugated siRNA duplex equivalents (µg/g) for comparability.

In the liver, unconjugated *Hprt* siRNA showed only minimal exposure (0.25 ± 0.07 µg/g at 10 mg/kg IV; Figure 7A and 7C), whereas Clone F-*Hprt* accumulated to 42.5 ± 6.9 µg/g, a ∼170-fold enhancement. This high level of accumulation was consistent with the strong *Hprt* mRNA knockdown observed in liver (Figure 6). A similar pattern was seen in the lung, where unconjugated siRNA reached only 0.063 ± 0.012 µg/g at 10 mg/kg compared with 1.53 ± 0.28 µg/g for Clone F-*Hprt* (Figure 7A and 7C), again aligning with the improved knockdown efficacy (Figure 6). In quadriceps, HELISA measurements revealed parental exposures of 0.188 ± 0.042 µg/g at 10 mg/kg compared to 1.11 ± 0.40 µg/g for Clone F-*Hprt* (Figure 7A and 7C), consistent with the robust gene silencing seen in muscle (Figure 6). In contrast, the kidney presented a different relationship between exposure and activity. LC-MS showed that unconjugated siRNA accumulated to 3.71 ± 0.30 µg/g at 10 mg/kg IV, higher than Clone F-*Hprt* (2.48 ± 0.57 µg/g; Figure 7A and 7C). Despite this relatively high exposure, Clone F-*Hprt* exhibited little or no mRNA knockdown in kidney (Figure 6). One possible explanation is that siRNAs accumulate in renal compartments with low HPRT expression, leading to limited functional engagement despite measurable bulk tissue concentrations.

To complement the quantitative tissue concentration measurements, we performed immunohistochemistry (IHC) for the target protein HPRT in liver and heart, following the procedures described in Materials and Methods, which showed a trend toward greater protein reduction with Clone F-*Hprt* treatment compared with unconjugated siRNA (Figure 7B). Together, these data indicate that tissue concentrations (Figure 7) align with tissue-specific activity (Figure 6) of Clone F-*Hprt* in liver, lung, and quadriceps, while kidney showed minimal activity.

Next, we analyzed the relative IGF1R levels in six peripheral tissues and the brain using western blot. IGF1R is composed of multiple subunits and is processed from a pro-IGF1R precursor. Multiple anti-IGF1R antibodies have been reported to yield both specific and non-specific bands in western blot (22). To distinguish the specific IGF1R bands, we compared the immunoreactivity of tissue lysates with that of MCF7 cell lysates and HeLa IGF1R knockout (KO) lysates (Figure 8A). Most tissue samples exhibited two prominent immunoreactive bands between ∼180-200 kDa and ∼80-110 kDa, which were absent in HeLa IGF1R KO lysates. We therefore considered these bands to be specific for IGF1R. Notably, lung and kidney lysates displayed additional smaller bands alongside the two major IGF1R-specific bands (Figure 8A).

**Figure 8.**
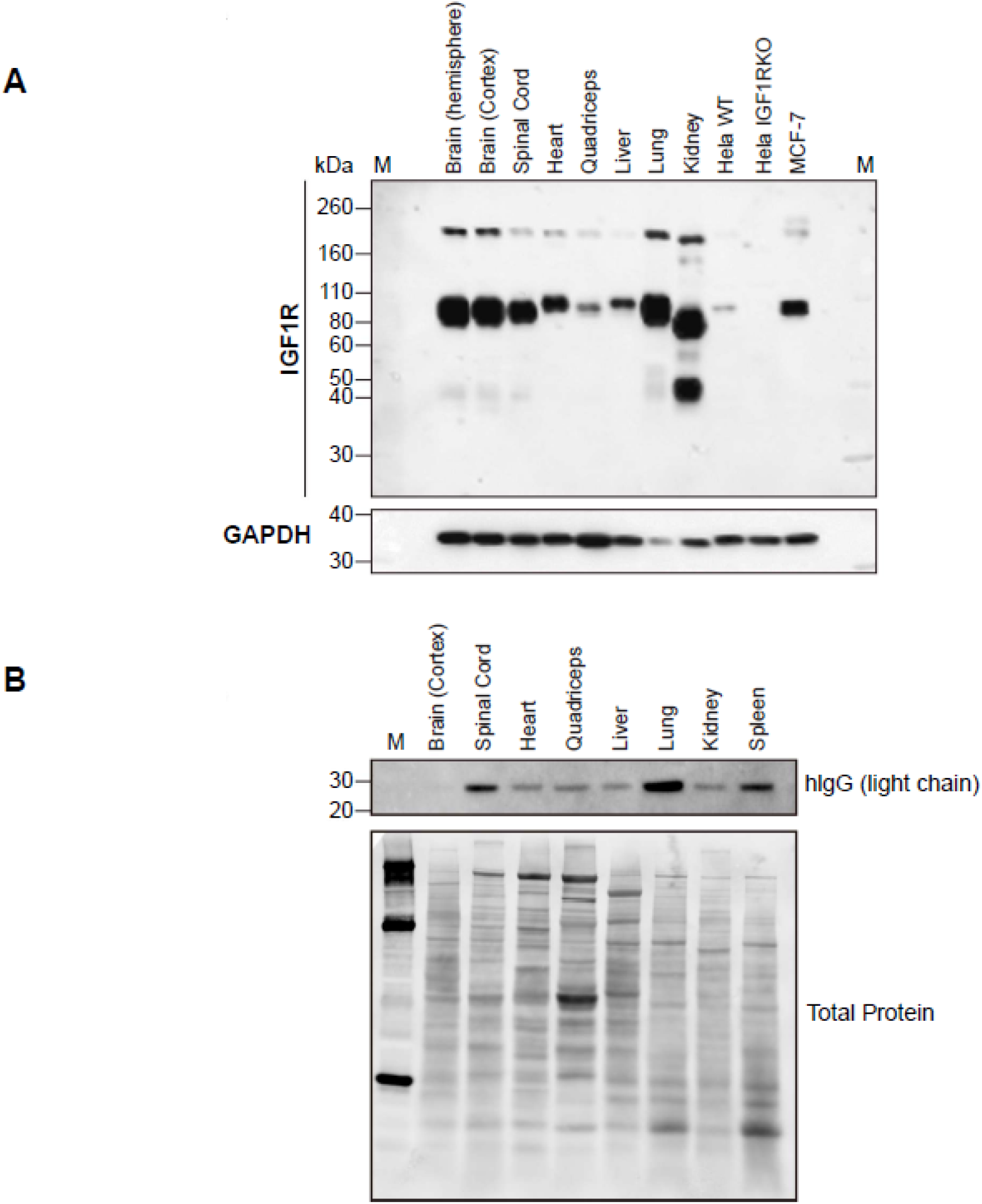
IGF1R expression and motavizumab-Clone F distribution in various mouse tissues.(A) Western blot analysis of IGF1R expression in brain and peripheral tissues, with MCF7 lysates as a positive control and HeLa IGF1R KO lysates as a negative control. Two major IGF1R-specific bands (∼180-200 kDa and ∼80-110 kDa) were observed in most tissues; additional smaller bands were detected in lung and kidney. (B) Western blot detection of hIgG light chain in tissues following IV administration of motavizumab-Clone F, showing a consistent band at 20–30 kDa across tissues. Total protein staining is included as a loading control.

To further evaluate the distribution of Clone F-*Hprt*, we assessed hIgG levels using an antibody against the hIgG light chain. Clear bands between 20-30 kDa were detected in all tissue lysates analyzed, with variable intensities across tissues (Figure 8B). We tested several loading controls and observed variability between tissues with all proteins examined (Figure 8A). As an alternative, total protein staining was used to visualize loading, which demonstrated overall similar protein levels across wells (Figure 8B).

Given 1) the global improvements in knockdown efficacy across brain regions, 2) the significantly higher efficacy observed in peripheral tissues following IV and SC dosing, and 3) the presence of IGF1R expression in both brain cells and peripheral tissues, we conclude that Clone F-*Hprt* delivery involves Clone F-mediated engaging with IGF1R facilitating transport into target cells.

## DISCUSSION

The brain protects itself from potentially hazardous materials outside the brain via a unique structural barrier called the blood–brain barrier (BBB). The BBB is formed by brain endothelial cells, tight junctions, pericytes, astrocytic endfeet, and basement membrane, covering 600-700 km of brain microvessels (23). This multiple structure generates a tight firewall with little fenestration, restricting entry of molecules >400 Da and favoring lipophilic compounds (24). Chemically modified, unconjugated siRNAs are hydrophilic and highly negatively charged, making brain penetration particularly challenging. Various approaches have been developed, and a few of them showed meaningful progress (6,25). However, methodologies currently used have their own pitfalls, for instance, toxicities of encapsulating materials in high dose (26) or complications with intrathecal delivery (27). In contrast, receptor-mediated transport (RMT) using antibody-oligonucleotide conjugates may provide a relatively safe and efficient route for systemic brain delivery.

Clone F-*Hprt*, given IV or SC, induced dose-dependent, significant knockdown in various brain regions. Significant reduction of *Hprt* mRNA in the cortex, striatum, thalamus, and cerebellum by Clone F-*Hprt* given SC contrasts with no or only minimal changes after IV administration of unconjugated siRNA. Notably, Clone F-*Hprt* also reduced *Hprt* in deeper brain regions such as thalamus and striatum, which are typically difficult to reach. The overall increased efficacy mediated by conjugation to the anti-IGF1R antibody shuttle in the CNS may be explained by several factors. First, Clone F-based shuttle may enhance penetration of siRNA into brain parenchyma through RMT. Elevated siRNA levels correlated with the regions showing higher efficacy, indicating more Clone F-*Hprt* reached the CNS. Since Clone F targets IGF1R expressed in brain endothelial cells and brain cells (14), it may have elevated cellular targeting once inside the brain. Consistent with our hypothesis, immunofluorescent images of brain sections from mice treated with Clone F-fused antibody exhibited clear cellular localization in addition to fluorescent signals along brain microvessels (Figure 4). In addition, conjugation of siRNA to an antibody increases molecular size, which may reduce clearance through the glymphatic system compared to unconjugated siRNA (28).

Extensive research has been conducted to develop effective delivery systems for siRNA targeting extrahepatic tissues. However, the number of ligand/receptor pairs capable of successfully delivering siRNA to these tissues remains limited, especially across the BBB. Most efforts have focused on gene modulation in muscle or tumors, with few successful approaches for CNS delivery (29-32). Interestingly, Clone F-*Hprt* induced potent, dose-dependent gene knockdown in multiple peripheral tissues, including liver, quadriceps, heart, and lung. The liver is known to be a highly perfused organ and a major site of non-RMT uptake so, the high potency of Clone F-*Hprt* in liver may follow a similar mechanism shown by others (31,32). Muscles, in contrast, presents a clear barrier to siRNA penetration (33). Given enriched expression of TfR, siRNA conjugates with anti-TfR antibodies have been reported as potential therapeutics for muscle diseases (31,32). IGF1R is also expressed in muscle according to our result and protein databases, although at lower level than brain (Figure 8). Since Clone F-*Hprt* retains the IgG structure despite the LALA effector-reducing mutation, its efficacy could involve not only IGF1R-mediated transport but also some degree of non-RMT uptake, as has been speculated for other antibody-oligonucleotide conjugates (31). However, consistent with findings in muscle, receptor-mediated delivery through IGF1R is likely the dominant and most productive mechanism in our system. Although kidney siRNA levels were measurable, knockdown remained minimal, indicating that most of the exposure reflects accumulation in compartments or cell types with poor target engagement, such as those with low *Hprt* expression. In addition, the large molecular size of Clone F-*Hprt* (∼200 kDa) likely limits filtration through the glomerular barrier, further reducing productive delivery. A similar non-productive exposure may also underlie the weak activity observed in spleen (5). Significant efficacy was also shown in heart and lung. Both tissues are highly segmented anatomically and physically. In particular, siRNA therapeutics targeting lung have had limited success with local administration, whereas Clone F-*Hprt* induced gene knockdown after systemic dosing, even at 1 mg/kg. Overall, these findings highlight Clone F-*Hprt* as a versatile platform for siRNA delivery. Certain neurological disorders, such as Rett syndrome, may benefit from combined CNS and peripheral intervention (34), and siRNA– Clone F conjugates may offer a unique advantage by providing broad efficacy across both compartments.

## DATA AVAILABILITIES

All data are incorporated into the article and its online supplementary data.

## ACKNOWLEDGEMENTS

We would like to thank qPCR core and histology core departments at Ionis, as well as Raul Alonzo for graphics support.

## Author contributions

Mengnan Tian (Conceptualization, Animal studies, Writing), Mehran Nikan (Conceptualization, Antibody-siRNA conjugation, Writing), Hien T. Zhao (Conceptualization, Project administration), Thazha P. Prakash (Conceptualization, Project administration), Stephanie Klein (Histology), John Matson (HELISA and LC-MS assays), Asa Wahlander (HELISA and LC-MS assays), Janel Huffman (HELISA and LC-MS assays), Michael Tanowitz (ELISA-based binding analysis), Sungwon An (Project administration, Writing), Jinwon Jung (Conceptualization, Project administration), Seung-Hwan Kwon (Methodology, Visualization, Data curation, Western blot), Miran Yoo (Methodology, Visualization, Data curation, Immunofluorescence), Dongin Kim (Methodology, Western blot), Sumin Hyeon (Methodology, Western blot), Weon-Kyoo You (Conceptualization, Methodology), Hakju Kwon (Conceptualization, Project administration), Sang Hoon Lee (Conceptualization, Project administration). All authors reviewed and edited the manuscript.

