## Supplementary information for "Systemic and Local Delivery of siRNA to the CNS and Periphery via Anti-IGF1R Antibody Conjugation"

^&^ Equal contributions

**List of contents**

**Figure S1	2**

Figure S2	2

Figure S3	2

Figure S4	3

Figures S5	4

Figures S6	5

Figures S7	6

Figures S8	6

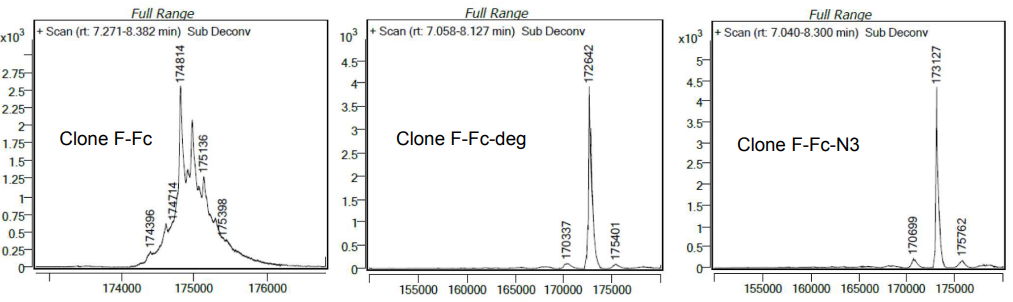

Figure S1. LC-MS spectra of Clone F-motavizumab, deglycosylated Clone F-motavizumab, and azide-functionalized Clone F-motavizumab are shown from left to right.

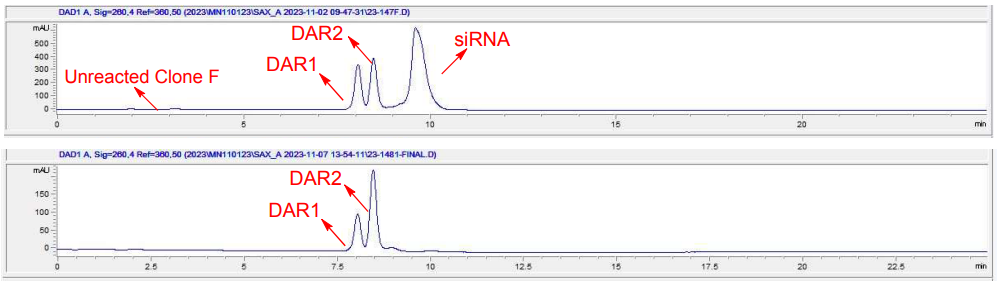

Figure S2. Anion-exchange chromatograms of the crude (top) and purified (bottom) Clone F-*Hprt* conjugate.

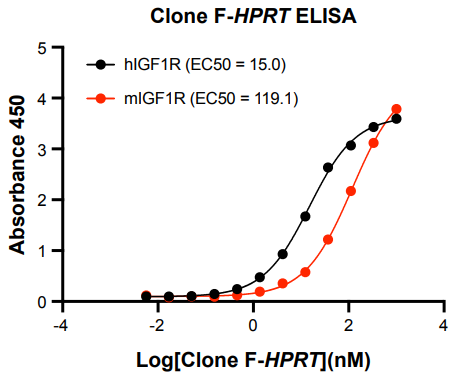

Figure S3. Binding affinities of Clone F-*Hprt* to human and mouse IGF1R ectodomain.

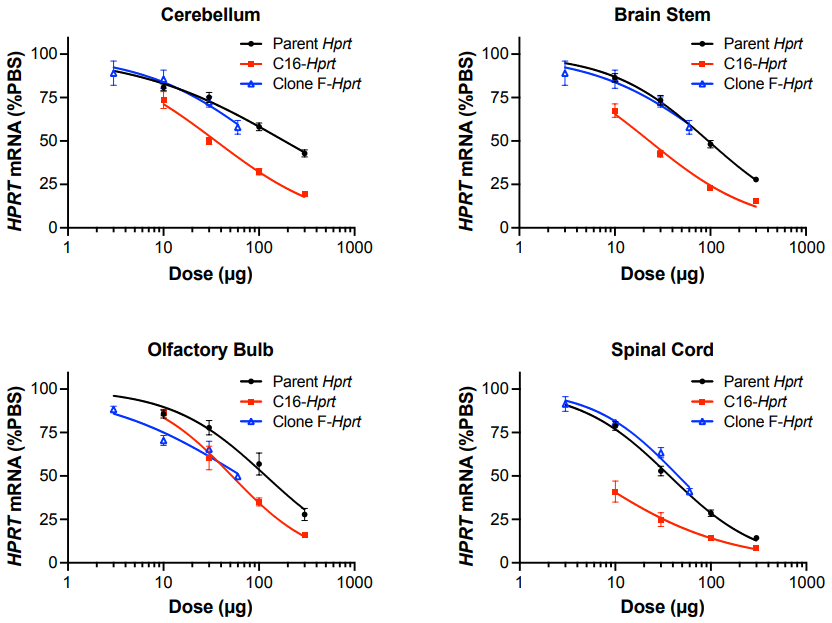

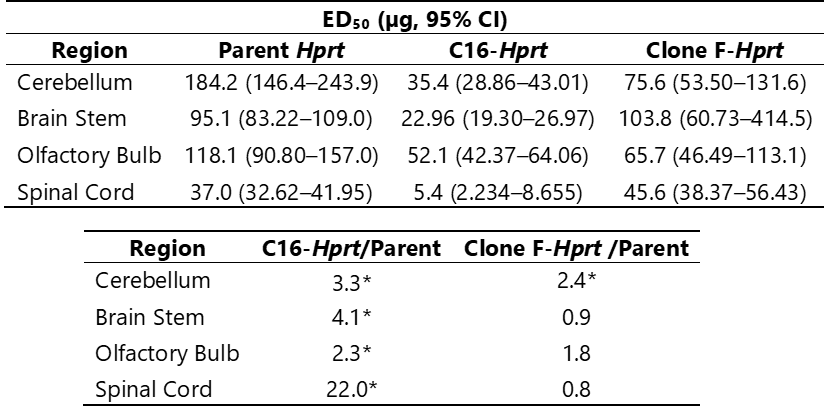

Figure S4. mRNA reduction across cerebellum, brain stem, olfactory bulb, and spinal cord after ICV administration of Clone F-*Hprt*, C16-*Hprt* or unconjugated *Hprt* siRNA. Each compound was administered at four different doses*. Hprt* mRNA levels were quantified by qRT-PCR from tissues collected two weeks post-dosing. Dose-response curves were generated using non-linear regression (log[agonist] vs. response) in GraphPad Prism, with the bottom constraint set to 0 and the top to 100. Data represent mean ± SEM. The unconjugated *Hprt* siRNA and C16-*Hprt* groups consisted of six animals per group (n = 6), while the Clone F-*Hprt* group consisted of four animals per group (n = 4). ED_50_s (95% CI) and fold-changes vs. the unconjugated siRNA are tabulated above. Asterisks denote a statistically significant difference (p < 0.05) in the potency (ED₅₀) of the conjugate compared to the unconjugated parent siRNA, as determined by statistical comparison of the dose-response curves.

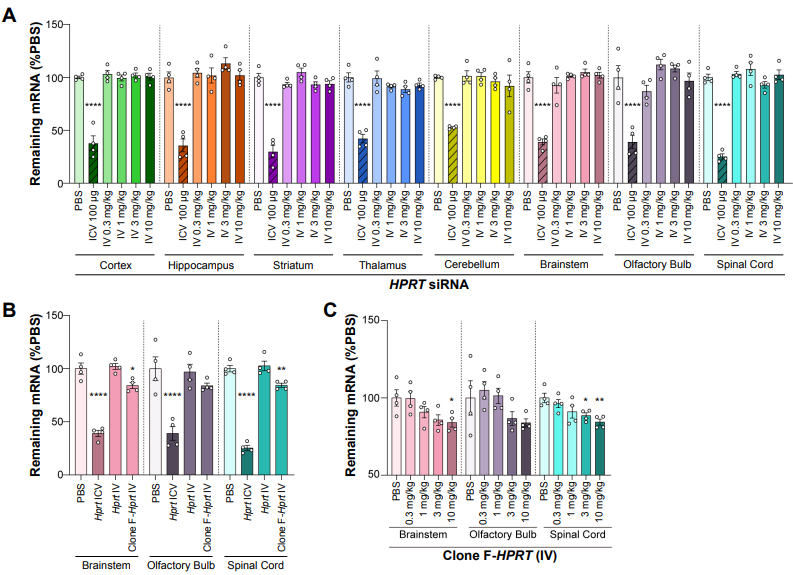

Figure S5. *Hprt* mRNA reduction in other CNS regions after peripheral administration. Mice received PBS, unconjugated *Hprt* siRNA, or Clone F-*Hprt* by IV injection (0.3-10 mg/kg, days 1 and 8); tissues were collected on day 15. A single ICV dose of 100 μg unconjugated siRNA was included as a positive control. *Hprt* mRNA levels were measured by qRT-PCR and expressed relative to PBS controls (mean ± SEM; n = 4). (A) No knockdown following IV administration of unconjugated *Hprt* siRNA (0.3-10 mg/kg) compared with ICV administration of unconjugated *Hprt* siRNA (100 µg) across different brain regions. (B) *Hprt* mRNA levels in brainstem, olfactory bulb, and spinal cord following ICV administration (100 μg), IV administration of unconjugated siRNA (10 mg/kg), or IV administration of Clone F-*Hprt* (10 mg/kg). (C) Dose-response analysis of Clone F-*Hprt* (0.3-10 mg/kg) in brainstem, olfactory bulb, and spinal cord. Modest knockdown was detected in the spinal cord at higher doses and in the brainstem at the highest dose, whereas the olfactory bulb showed only non-significant trends. Significance was determined by one-way ANOVA with Dunnett’s multiple comparisons test (*p < 0.05; **p < 0.01; ***p < 0.005; ****p < 0.001).

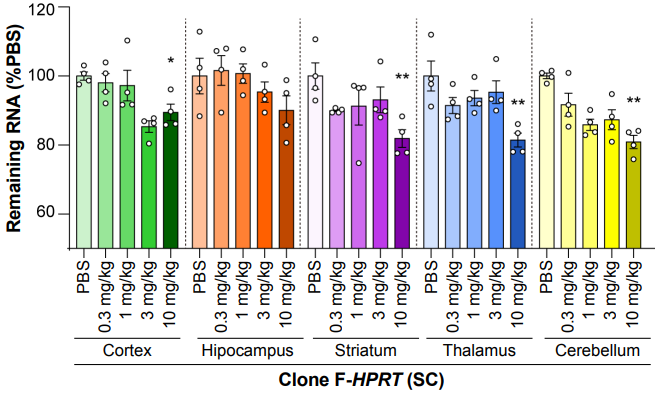

Figure S6. Dose-dependent knockdown of *Hprt* mRNA across CNS regions after SC administration of Clone F-*Hprt*. Animals were dosed SC with Clone F-*Hprt* (0.3-10 mg/kg, days 1 and 8); tissues were collected on day 15. *Hprt* mRNA levels were measured by qRT-PCR and expressed relative to PBS controls (mean ± SEM; n = 4). Significance was determined by one-way ANOVA with Dunnett’s multiple comparisons test (*p < 0.05; **p < 0.01; ***p < 0.005; ****p < 0.001).

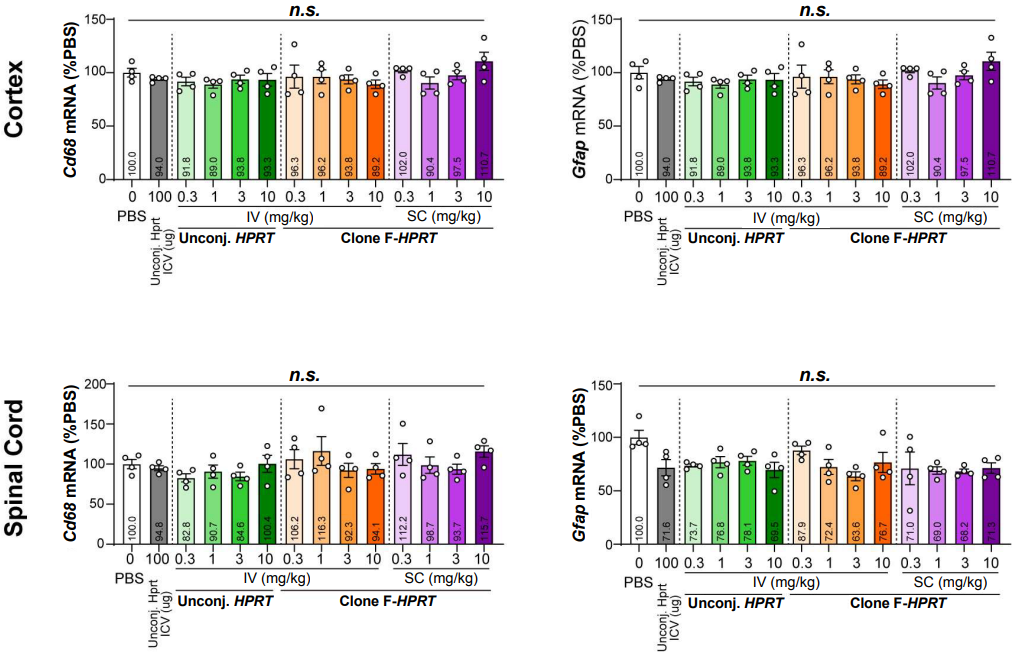

Figure S7. No significant alterations of brain Cd68 and Gfap mRNA levels after Clone F-*Hprt* administration. Statistical analysis showed no significant differences between groups (N = 4). n.s.= not significant.

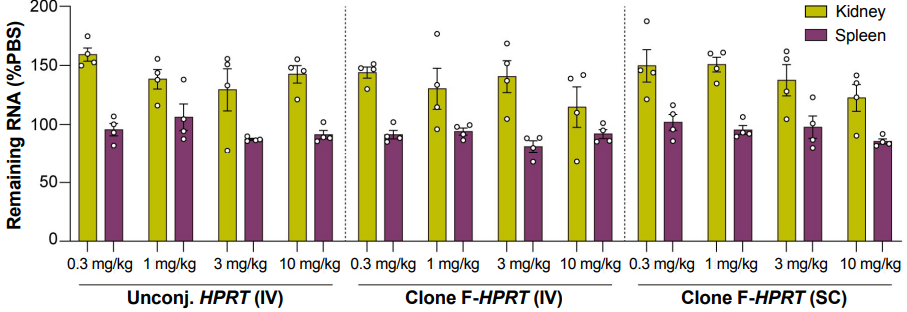

Figure S8. *Hprt* mRNA levels in kidney and spleen following escalating doses of unconjugated *Hprt* siRNA (IV) or Clone F-*Hprt* (IV and SC), showing limited to no target engagement.
